# Two disparity channels in human visual cortex with different contrast and blur sensitivity

**DOI:** 10.1101/2023.09.14.557625

**Authors:** Milena Kaestner, Yulan D. Chen, Caroline Clement, Alex Hodges, Anthony M. Norcia

**Author notes:** Conflicts of interest: None to declare.

## Abstract

**Purpose:** Our goal is to describe the contrast and blur sensitivity of multiple horizontal disparity sub-systems and to relate them to the contrast and spatial sensitivities of their monocular inputs.

**Methods:** Steady-State Visual Evoked Potentials (SSVEPs) amplitudes were recorded in response to Dynamic Random Dot Stereograms (DRDS) alternating at 2 Hz between zero disparity and varying magnitudes of crossed disparity for disparity plane and disparity grating stimuli. Half-image contrasts ranged between 2.5 and 80% and over a range of Gaussian blurs from 1.4 to 12 arcmin. Separate experiments measured contrast and blur sensitivity for the monocular half-images.

**Results:** The first and second harmonics disparity responses were maximal for disparity gratings and for the disparity plane condition, respectively. The first harmonic of the disparity grating response was more affected by both contrast and blur than was the second harmonic of the disparity plane response which had higher contrast sensitivity than the first harmonic.

**Conclusion:** The corrugation frequency, contrast and blur tuning of the first harmonic suggest that it reflects activity of neurons tuned to higher luminance spatial frequencies that are selective for relative disparity, whereas the second harmonic reflects the activity of neurons sensitive to absolute disparity that are driven by low monocular spatial spatial frequencies.

**Translational Relevance:** SSVEPs to DRDS provide two objective neural measures of disparity processing, the first harmonic – whose stimulus preferences are similar to those of behavioral stereoacuity – and the second harmonic that represents an independent disparity-specific, but not necessarily stereoscopic mechanism.

## INTRODUCTION

Stereo-acuity, our ability to sense depth differences provided by horizontal disparity cues, is among the finest discriminations made by the visual system. Stereo thresholds under optimal conditions are under 10 seconds of arc (McKee, 1983), allowing depth discrimination up to a range of over a kilometer (Allison et al., 2009). Psychophysical studies have suggested that stereoacuity is subserved by temporally sustained processing mechanisms (Ogle and Weil, 1958; Harwerth and Rawlings, 1977; Westheimer and Pettet, 1990) that rely on high spatial frequency image content (Schmidt, 1994; Hess et al., 1999). Stereo acuity is variable among the general population, exhibiting a distribution of acuities that is skewed with a long tail at higher stereo-thresholds (Coutant and Westheimer, 1993; Bosten et al., 2015; Chopin et al., 2019). Importantly, stereopsis is strongly affected by strabismus (eye-misalignment) and/or anisometropia (unequal refractive error in the two eyes) present during early development and is thus an important metric for clinical studies of binocular function.

In clinical applications, many previous studies have used variations on random-dot stereograms in children who are able to respond to instructions and provide a verbal or manual response (Simons, 1981; Heron et al., 1985; Oduntan et al., 1998; Tomac and Altay, 2000; Kulp and Mitchell, 2005; Birch et al., 2008; Anketell et al., 2013; Drover et al., 2017; Baldev et al., 2022). It would be desirable to establish an objective index of stereoacuity for pre-verbal children who are the target of clinical intervention. To this end, there have been several studies using the preferential looking technique to measure random-dot stereoacuity in infants (Birch and Salomao, 1998; Birch et al., 2005; Morale et al., 2021). Visual evoked potentials (VEPs) can also be used to estimate the stereoacuity of infants (Birch and Petrig, 1996; Norcia et al., 2017). These VEP studies have used a variant of the random dot stereogram (RDS), the dynamic RDS (DRDS), to evoke cortical responses to changes in binocular disparity. The temporally uncorrelated frames of a DRDS obscure monocular motion cues associated with changes in disparity.

Early work with DRDSs assumed that an evoked response to a DRDS was sufficient to demonstrate the presence of stereopsis (Julesz et al., 1980). However, stereopsis is a *sense* of depth from horizontal disparity and work in macaque (Cumming and Parker, 1997) has suggested that simply detecting disparity is not a sufficient criterion for stereopsis. Their argument was based on the finding that anti-correlated RDSs evoke a disparity-tuned cell responses in macaque V1, even when these stereograms do not give rise to a percept of depth.

Moreover, recent work using VEPs has suggested that young infants have a comparable response to horizontal and vertical disparities, unlike adults who have greater sensitivity for horizontal disparity than for vertical disparity (Norcia et al., 2017). Vertical disparities also do not give rise to a percept of depth, so a stronger criterion for functional stereopsis is needed than the simple presence of an evoked response to changing disparity.

What might such a criterion be for evoked responses? The standard psychophysical probe of stereopsis is the discrimination of the sign of disparity, rather than the mere detection of the presence or absence of disparity. Recent work has suggested that the direction of motion in depth can be decoded from high-density VEP recordings (Himmelberg et al., 2020). However, it is unclear whether this approach could be used to measure stereoacuity, as the results were obtained by recording many trials with highly supra-threshold disparities.

Previous VEP studies have used other approaches to linking parameters of the evoked response to behavioral disparity sensitivity. Norcia and colleagues (Norcia et al., 1985a) compared psychophysical disparity thresholds to VEP thresholds and found them to be similar (see also (Wesemann et al., 1987; Kohler et al., 2018). Kaestner and co-workers (Kaestner et al., 2022) showed that the first harmonic response to the onset and offset of a disparity grating shares the same tuning for cyclopean spatial frequency as does psychophysics. They also found that the simultaneously recorded second harmonic was untuned for cyclopean spatial frequency – unlike perception, and this suggested that the VEP signal is comprised of a mixture of responses to absolute and relative disparity, with the 1^st^ harmonic largely reflecting the latter and the 2^nd^ harmonic largely the former.

Here we use three additional stimulus manipulations that are known to affect perceptual stereo-acuity to validate VEP response components as measures of stereopsis: the effect of disparity references as in Kaestner et al., (2022) and reductions of the contrast or high spatial frequency content in the half-images. These manipulations are motivated by prior work indicating that stereoacuity is strongly dependent on disparity references (Westheimer, 1979; McKee et al., 1990; Kumar and Glaser, 1991; Andrews et al., 2001), image contrast (Halpern and Blake, 1988; Legge and Gu, 1989; Westheimer and Pettet, 1990; Cormack et al., 1991) and blur (Brooks et al., 1996; Atchison et al., 2020a; Atchison et al., 2020b). Contrast reductions affect all spatial frequencies in the monocular half-images and are thus somewhat non-specific. Spatial blur provides a more specific manipulation of the spatial inputs to stereopsis as it selectively removes high spatial frequency monocular inputs. We also used these manipulations to further characterize putative relative– and absolute-disparity sensitive mechanisms reflected in the 1^st^ and 2^nd^ response harmonics.

Here we find that the first harmonic of the changing disparity response to disparity gratings shares several properties associated with psychophysical stereopsis: thresholds are in the hyper-acuity range, are dependent on references, and are highly dependent on the contrast and high spatial frequency content of the half-images. The second harmonic response to disparity planes, by contrast has a higher disparity threshold, a lower contrast threshold, is less dependent on references, and is less dependent on contrast and blur, the latter suggesting that it derives from mechanisms that use low spatial frequency portions of the monocular spatial frequency spectrum.

## METHODS

### Participants

All participants were recruited from the Stanford University community and were screened for normal or corrected-to-normal vision, ocular diseases, and neurological conditions. Visual acuity was measured using a LogMAR chart (Precision Vision, Woodstock, IL, USA) and was better than 0.1 LogMAR in each eye, with less than 0.3 LogMAR acuity difference between the eyes. Stereoacuity was measured with the RANDOT stereoacuity test (Stereo Optical Company, Inc., Chicago, IL, USA) with a pass score of 50 arcsec or better.

A total of 44 unique participants meeting these inclusion criteria contributed data to the five experiments reported here. Data from an additional 9 participants were excluded due to a lack of signal at the 20 Hz dot-update rate or at the harmonics of the 2 Hz disparity update rate. Because participants partially, but not fully overlapped across recording protocols, sample sizes, age and sex demographics are presented separately below for each experiment. Informed written and verbal consent was obtained from all participants prior to participation under a protocol approved by the Institutional Review Board of Stanford University.

### Visual Display and DRDS Stimuli

We first describe the common features of the DRDS stimuli that were used in all experimental protocols. Individual protocols, described separately below, differed in terms of their cyclopean (depth corrugation) spatial frequencies, half-image contrasts, and levels of half-image blur.

All stimuli were displayed on a SeeFront 32” autostereoscopic 3D monitor running at a refresh rate of 60 Hz. The SeeFront display comprises a TFT LCD panel with an integrated lenticular system that interdigitates separate images for the left and right eyes on alternate columns of the 3840 x 3160 native display resolution. In 3D mode, the effective resolution in 1920 x 1080 pixels per eye. Mean luminance was 111.4 cd/m^2^ (min-max range 2 – 220 cd/m^2^) as requested by the stimulus generation software after in-house calibration and gamma linearization. The viewing distance was 70 cm which is within the optimal range for the adult 65 mm average inter-pupillary distance, per the manufacturer’s specifications. At this distance, the monitor subtended. 53.7° x 31.8° with the stimulus display area being 31.8° x 31.8°. The SeeFront device monitors the participants’ head position via an integrated pupil location tracker and shifts the two eyes’ views to compensate for head motion, thus ensuring that half-images are projected separately into each eye. Head positioning was checked periodically for each participant by asking them to report on the separate visibility of the nonius lines.

The stimulus frames comprised dynamic random-dot stereograms (DRDS) generated in MATLAB using Psychtoolbox-3 (Brainard, 1997; Pelli, 1997; Kleiner et al., 2007). These frames were presented via a custom Objective C application with minimal jitter or frame dropping. Dot positions in the DRDS updated at a rate of 20 Hz. Random dots were presented within a circular aperture (29.4° diameter) embedded within a square 31.8° by 31.8° 1/f noise fusion lock that was used to stabilize eye gaze and vergence angle (see Figure 1a). The fusion lock was at zero disparity. A 1.2° wide ring of binocularly uncorrelated dots was placed between the edge of the fusion lock and the DRDS to reduce the availability of relative disparity cues arising from the edge of the zero-disparity fusion lock and the DRDS (Cottereau et al., 2012a). These uncorrelated dots were identical to the stimulus dots in size (6 arcmin) and density (15 dots/degree^2^) but were always shown at full contrast (100% Michelson) and their positions did not update. The visible diameter of the DRDS stimulus was 27°. To further control the eye position of participants, and to aid stable binocular fusion, nonius lines were placed at central fixation where the length of each line was 1° with 0.3° separation between upper and lower lines.

**Figure 1:**
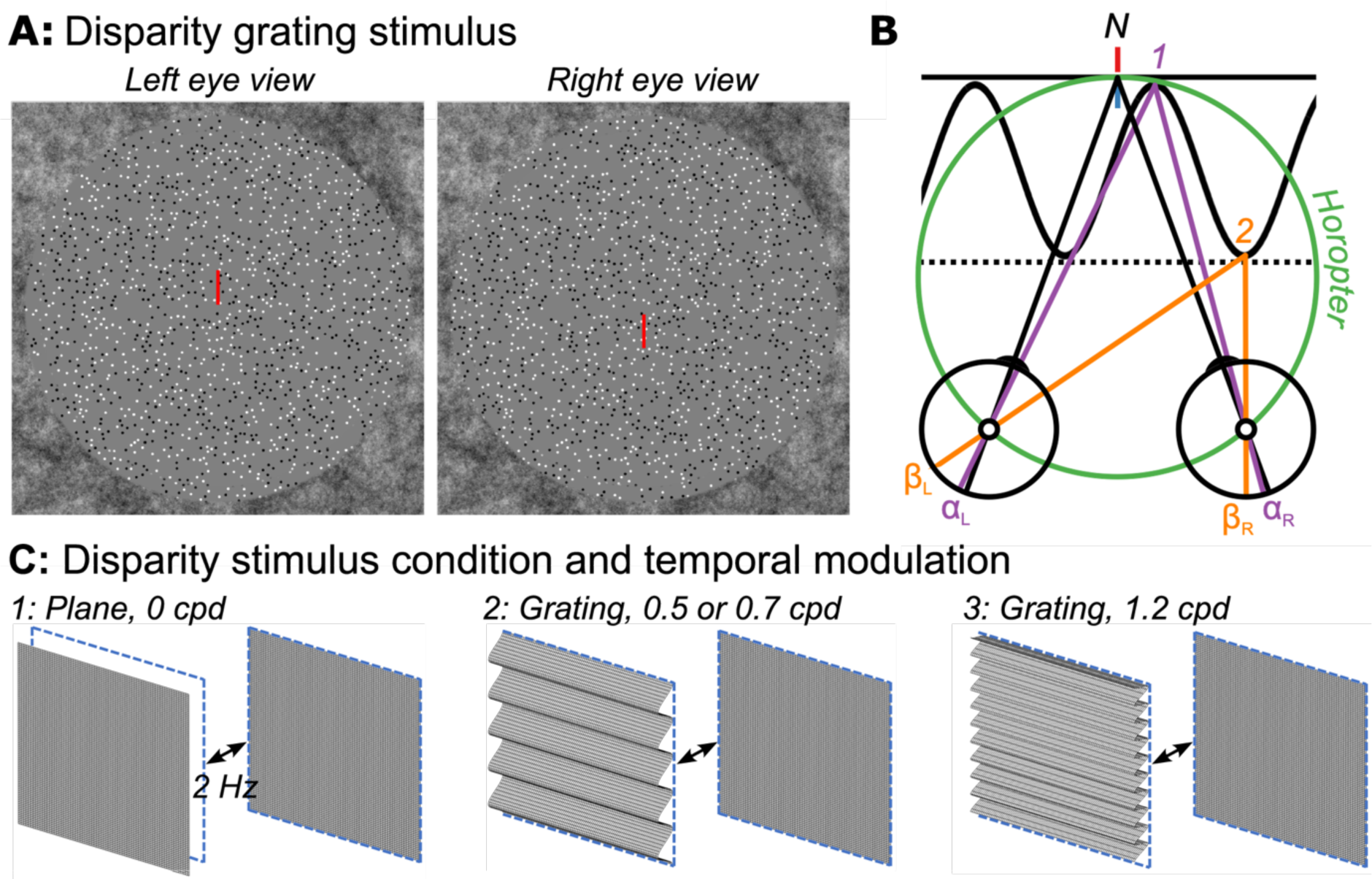
**a:** Random-dot stereopair used in the main experiment depicting a sinusoidal disparity grating (crossed disparity when cross-fused). The dots in the actual experiment were dynamic. **b:** Top-down schematic illustration of disparity plane (absolute) and disparity grating (relative plus absolute disparity) stimuli. Fixation point N (nonius lines) is on the zero-disparity plane defined by the horopter (green Vieth-Muller circle). Point 1 (purple) is also on the horopter and angles α_L_ and α_R_ are equal, meaning that the absolute disparity given by α_L_ – α_R_ is zero. Point 2 (orange) is either on a second, disparate plane (dotted line) or on the peak of a disparity grating (sinusoidal line). Here the absolute disparity, β_L_ – β_R_, is non-zero. The relative disparity is the difference between the two absolute disparities, (α_L_ – α_R_) − (β_L_ – β_R_), and varies depending on the location on the sinewave but its magnitude is independent of fixation. Disparity along the second plane (dotted line) is constant at a non-zero value. **c:** Schematic of disparity plane stimulus (1) that involves the temporal modulation at 2 Hz between crossed disparity, ‘disparity on’, and zero disparity, ‘disparity off’ phases. Schematic illustration of disparity grating stimuli (2, 3) that involve 2 Hz temporal modulation between a cyclopean grating portrayed with crossed disparity and a zero-disparity plane. Participants were asked to detect a brief color change on the nonius fixation lines.

### Contrast and blur manipulations

In Experiments 1-4, we measured VEP amplitude as a function of disparity at a series of half-image contrasts and various amounts of half-image blur in different recording protocols to be described in detail below. The disparity profiles we used are illustrated schematically in Figure 1b which illustrates disparity plane modulation and disparity grating modulation conditions in top-down view. In each case, disparity modulation is relative to the zero-disparity plane defined with reference to the horopter (green circle and nonius lines). In the disparity plane condition, the stimulus alternates between zero disparity (solid black line, label 1) and varying amounts of crossed disparity (dotted line; label 2). This stimulus varied in absolute disparity with minimized contributions of relative disparity due to the uncorrelated dot region between the active display and surrounding zero disparity fusion lock. In the disparity grating conditions, the display alternated between zero disparity and a crossed disparity corrugated surface of varying disparities. This display contains both absolute and relative disparities (see legend for geometric definitions of absolute and relative disparities). We used three cyclopean spatial scales of the disparity grating modulation (corrugation frequencies), 0.5, 0.7 and 1.2 cpd over different protocols, as described below.

Half-image contrast was defined on the Michelson definition in the space domain and blur was generated by convolving the half-images with a Gaussian blur kernel using the MATLAB ‘imgaussfilt’ function. The Gaussian blur kernel was defined by its standard deviation, given to the function in units of pixels. For figures and analysis, the pixel unit was converted to arcmin given the viewing distance of the participant and the pixel resolution of the half-images. To illustrate the effects of the contrast and blur manipulations on the half-images, Figure 2 shows example frames at different contrasts (top) and blur levels (bottom), along with their Fourier power spectra. Varying contrast shifts the entire response spectrum vertically on the plot, while varying blur progressively decreases the high spatial frequency image content, leaving the lowest spatial frequency power largely unaffected.

**Figure 2.**
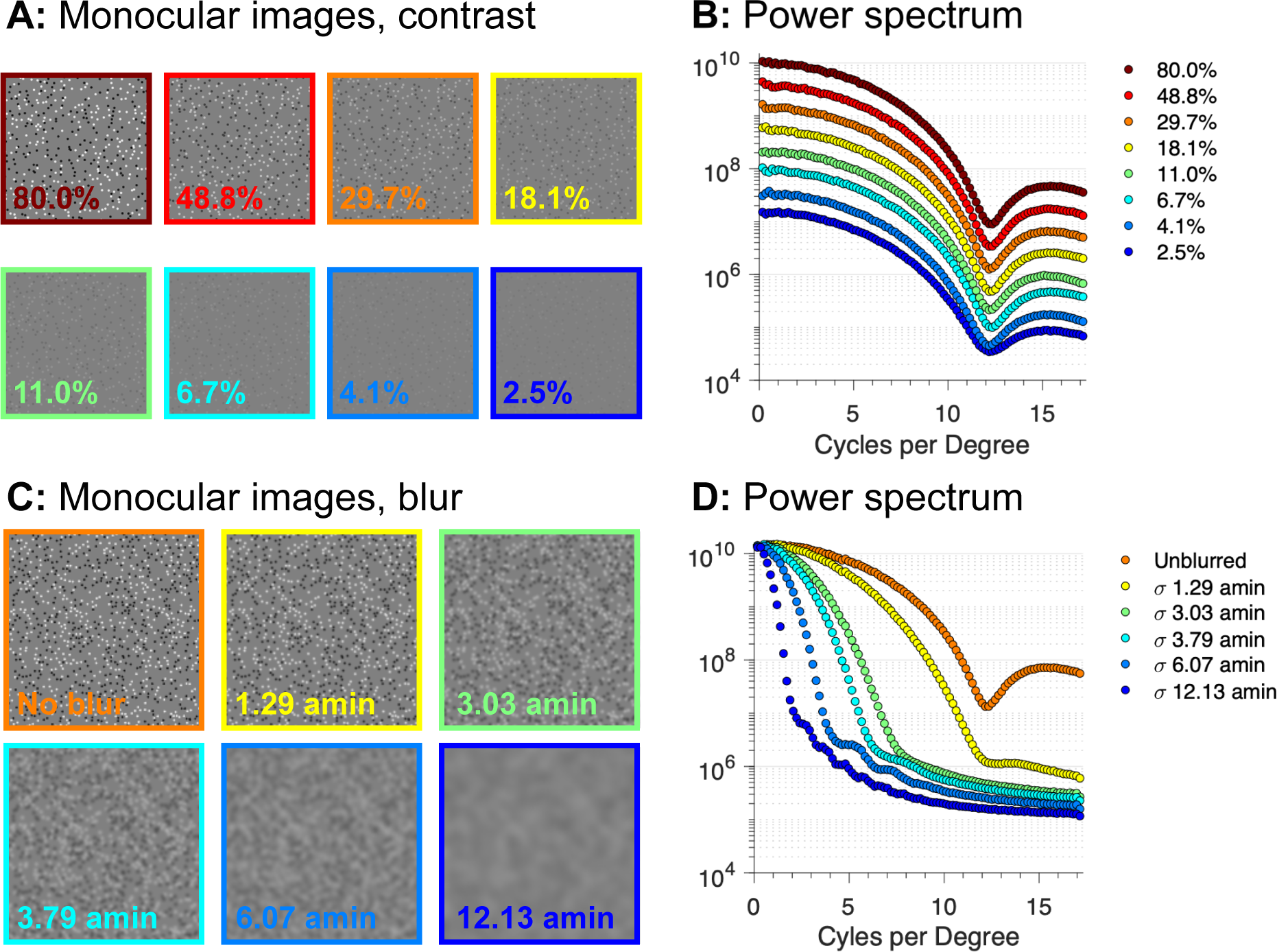
Example half-images for contrast variation (top left) and blur variation (bottom left) with corresponding Fourier power spectra. Contrast in the example images is specified on the Michelson definition. The blur levels are specified in terms of the standard deviation of the gaussian kernel in terms of visual angle (arc min). Contrast reduction affects all spatial frequencies while blur manipulation preferentially affects high spatial frequencies.

### Swept parameter paradigms

For all experiments and stimulus types, response functions were measured using swept-parameter protocols in which the visibility of the stimulus parameter of interest was “swept” from invisible to visible over 10 equal log-spaced steps. In Experiments 1-4, the swept parameter was disparity and the different stimulus conditions varied in half-image contrast (Experiments 1 and 2) or blur level (Experiments 3 and 4). In Experiment 5, the swept parameter was either contrast or blur, with the disparity being zero. This experiment established limits on the visibility of the dynamic dots in the half-images.

Individual sweep trials began with a 1 s prelude in which the display presented the first 60 frames of the upcoming sweep, allowing the stimulus onset transient to dissipate and the adaptive filter (see below) to reach a steady state. This prelude was followed seamlessly by a 10 s stimulus presentation period, during which disparity, contrast or blur amplitude was the swept parameter. The trial ended with a 1 s postlude, recycling the last 60 frames of the stimulus. There was a minimum of 2 s mean luminance interval between subsequent trails, during which participants were instructed to blink as needed.

During stimulus trials, participants were asked to attend to the nonius lines and to press a button to indicate when a color change occurred. The purpose of the task was to encourage fixation at the centre of the active display area, with convergence on the plane of the display, and to monitor the vigilance state of the participants. The initial duration of the colour change was 0.5 s and was varied on a staircase that maintained an 82% correct level of performance.

### Experiments 1 and 2: Contrast dependence of plane and grating disparity responses

We measured the contrast dependence of the changing disparity response in two experimental protocols using partially overlapping participants. Experiment 1 (N=8, 4 male, mean age = 28.1 years, 3 EEG sessions per participant) compared the disparity response function for a disparity plane and a 0.7 cpd disparity grating, and Experiment 2 (N=8, 2 male, mean age = 29.5 years, 3 EEG sessions per participant) compared the disparity response function for 0.7 and 1.2 cpd disparity gratings. In both protocols, the contrast of the dots in the DRDS was set at eight different levels in each of the different stimulus conditions, with contrast ranging in equal log steps from 2.5 – 80%.

To measure the disparity response function, peak-to-peak disparity amplitude of the grating or plane was “swept” in 10 equal log steps over each 10 s stimulus presentation. The stimulus completed two disparate/non-disparate cycles at each 1 sec step in the disparity sweep. Because disparity sensitivity varies as a function of cyclopean spatial frequency, different sweep ranges were chosen for plane and grating stimuli. For the disparity plane condition, disparity amplitude was swept between 0.5 and 8 arcmin, whereas for disparity grating conditions the disparity amplitude was swept between 0.2 and 6 arcmin for the 0.7 cpd grating and between 0.5 and 8 arcmin for the 1.2 cpd grating. Optimal sweep ranges were chosen based on pilot experiments and the results of a previous experiment (Kaestner et al., 2022), such that the disparity response emerged from the noise in the first half of the sweep and did not saturate towards the end.

In each contrast protocol, there were 16 conditions total – 8 contrast levels for each disparity type. Blocks of 10 trials per condition were recorded in 3 separate sessions for a total of 30 trials per condition per participant, *e.*g. 300 sec of data over the 10 levels of the sweep with 30 sec of data per level, per participant. The order of blocks was randomised between participants and breaks were permitted between each block.

### Experiments 3 and 4: Blur dependence of plane and grating disparity responses

In Experiments 3 and 4, response functions were measured under varying levels of blur applied equally to each of the half-images. As for the contrast experiments, blocks of 10 trials were run at a given blur level and the order of blocks was randomized across participants. In Experiment 3 (N=15; average age 24, 8 female, one EEG session per participant) disparity response functions were measured for disparity plane, 0.5 cpd and 1.2 cpd grating conditions with 0, 1.37 and 4.1 arcmin of blur, using sweep ranges of 0.5 to 8 arcmin for the plane and 1.2 cpd grating and a sweep range of 0.2 to 6 arcmin for the 0.5 cpd disparity grating.

In Experiment 4 (N=15; average age 26, 10 female, one EEG session per participant) disparity response functions were measured with a larger range of blur values. Responses to the disparity plane were measured with 0, 6.07 and 12.13 arcmin of blur using sweep ranges of 0.5-8, 0.75-12, and 0.75-12 arcmin disparity, respectively. Responses to an 0.5 cpd disparity grating were measured at 0, 3.03 and 6.07 arcmin of blur using sweep ranges of 0.2 to 6, 0.2 to 6 and 0.5 to 8 arcmin disparity. Responses to a 1.2 cpd disparity grating were measured at 0, 3.03 and 6.07 arcmin of blur using sweep ranges of 0.5 to 8, 0.75-12 and 0.75-12 arcmin disparity. Sweep ranges were shifted between conditions to optimally capture the variation in the response amplitude across different blur levels.

### Experiment 5: Effects of contrast and blur on monocular half-image visibility

Experiment 5 measured contrast and blur thresholds for the monocular half-image content (N=22, average age 25.3, 12 female, one EEG session per participant). The base display comprised a zero disparity DRDS updating at 20 Hz. Contrast thresholds were measured by sweeping the half-image contrast over 10 equal log steps between 1.25 and 40% contrast. These steps of contrast were alternated at 2 Hz with 0% contrast half-images presented within the central 29.5 deg disk region of the display. A blur threshold was measured by modulating the magnitude of gaussian blur imposed on a nominally 100% contrast unblurred DRDS within the same disk region. The images alternated at 2 Hz between blurred and unblurred. The magnitude of blur was swept between 0.3 and 7.6 arcmin. A total of 20 10 sec sweep trials were collected for the contrast and blur conditions (*e.g.* 200 sec data per condition).

### EEG Acquisition and Pre-Processing

High-density, 128-channel electroencephalograms (EEG) were recorded using HydroCell electrode arrays and an Electrical Geodesics Net Amps 400 (Electrical Geodesics, Inc., Eugene, OR, USA) amplifier. The EEG was sampled natively at 500 Hz and then resampled at 420 Hz, giving 7 data samples per video frame. The display software provided a digital trigger indicating the start of the trial with millisecond accuracy. The data were filtered using a 0.3 – 50 Hz bandpass filter upon export of the data to custom signal processing software. Artifact rejection was performed in two steps. First, the continuous filtered data were evaluated according to a sample-by-sample thresholding procedure to locate consistently noisy sensors. These channels were replaced by the average of their six nearest spatial neighbours. Once noisy channels were interpolated in this fashion, the EEG was re-referenced from the Cz reference used during the recording to the common average of all sensors. Finally, 1 sec EEG epochs that contained a large percentage of data samples exceeding threshold (30 – 80 microvolts) were excluded on a sensor-by-sensor basis.

### Fourier Decomposition via Recursive Least Squares Filtering

The steady-state VEP (SSVEP) amplitude and phase at the first four harmonics of the disparity update frequency (2 Hz) were calculated by a Recursive Least Squares (RLS) adaptive filter (Tang and Norcia, 1995). The RLS filter consisted of two weights – one for the imaginary and the other for the real coefficient of each frequency of interest. Weights were adjusted to minimise the squared estimation error between the reference and the recorded signal. The memory-length of the filter was 1 s, such that the learned coefficients were averaged over an exponential forgetting function that was equivalent to the duration of one bin of the disparity sweep. Background EEG levels during the recording were derived from the same analysis and were calculated at frequencies 1 Hz above and below the response frequency, e.g., at 1 and 3 Hz for the 2 Hz fundamental. Finally, the Hotelling’s T^2^ statistic (Victor and Mast, 1991) was used to test whether the VEP response was significantly different from zero. These measurements were used as part of the threshold estimation algorithm (see below).

### Dimension Reduction via Reliable Component Analysis

Reliable Components Analysis (RCA) was used to reduce the dimensionality of the sensor data into interpretable, physiologically plausible linear components (Dmochowski et al., 2015). This technique optimizes the weighting of individual electrodes to maximize trial-to-trial consistency of the phase-locked SSVEPs. Components were learned on RLS-filtered complex value data, and were learned on the 1F1, 2F1, 3F1 and 4F1 responses across all trials and all participants. Due to differences in topography for absolute and relative disparity mechanisms, we learned separate components for plane and grating conditions, the latter pooling across 0.5, 0.7 or 1.2 grating conditions on a per experiment basis. The Rayleigh quotient of the cross-trial covariance matrix divided by the within-trial covariance matrix was decomposed into a small number of maximally reliable components by solving a generalized eigenvalue problem. Each component can be visualized as a topographic map by weighting the filter coefficients by a forward model (Haufe et al., 2014; Dmochowski et al., 2015) and yields a complex-valued response spectrum for that component.

Participant-level sensor-space data for the dominant 1F1 and 2F1 harmonics were spatially filtered through the weights of the most reliable spatial filter, RC1. These filtered, participant-level amplitude and phase estimates for signal (1F1 and 2F1) and noise (side bands of the 1F1 and 2F1 harmonics, respectively) frequencies were calculated by taking the vector mean across real and imaginary components, across all trials within the same condition. Amplitude was calculated by taking the square root of the sum of the squared real and the squared imaginary components. Phase was calculated by taking the inverse tangent of the real and imaginary components.

### Group-level Disparity Response Functions

Group-level response functions at 1F1 and 2F1 after spatial filtering via RCA were estimated by determining the magnitude of the projection of each participant’s response vector on to the group vector average (Hou et al., 2009) for each level of the disparity, contrast or blur sweeps. Each individual participant’s response vector amplitude was multiplied by the cosine of the phase difference between it and the group mean vector (Hou et al., 2009). The scalar magnitude of these projections was then used to calculate the group mean projected amplitude and its standard error.

### Disparity, contrast and blur threshold estimation

Sensory thresholds for disparity, contrast and blur were estimated from by extrapolating the group average amplitude *vs* stimulus parameter function to zero amplitude. Thresholds were estimated at the first (1F1) and second (2F1) harmonics of the parameter’s modulation frequency. Details of the extrapolation algorithm are described elsewhere (Kaestner et al., 2022). Briefly, the algorithm sought sections of the response function that were above the noise level that could be well-approximately by a linear fit. The range for the extrapolation included the longest monotonically increasing portion of the response function, starting from when the signal began to rise out of the experimental noise level determined from adjacent EEG frequencies. The x-axis intercept was taken as the neural threshold – the disparity, contrast or blur at which the cortical response would have been zero in the absence of additive EEG noise. Errors on the threshold estimate were derived via the Jackknife direct estimation method (Miller, 1974; Norcia et al., 1985b; Miller et al., 1998).

## RESULTS

We measured SSVEP responses to stimuli containing absolute disparity modulation (the disparity plane stimulus) and a combination of absolute and relative disparity modulation (the disparity grating stimuli) as a function of disparity amplitude, disparity spatial frequency, Michelson contrast and blur levels. We also measured responses to modulation of blur and contrast to establish visibility limits for the half-images. As 1F1 and 2F1 responses were consistently measurable in all experimental conditions, the analysis of the evoked responses focusses on these two response harmonics.

RCA was used to derive the maximally reliable response component from the feature matrix comprised of electrodes and harmonics. Figure 3 shows the spatial forward model projections of the RCA weight distributions (response topographies) obtained for our different disparity modulation conditions for the first, maximally reliable component, RC1. These topographies are maximal over midline electrodes anterior to Oz and are broadly similar to each other and to previous measurements with similar stimuli (Norcia et al., 2017; Kaestner et al., 2022). Higher-order RCs were dissimilar across stimulus types or comprised non-significant signals, and thus were not readily comparable. Analysis was carried out by projecting data through RC1 learned for each condition.

**Figure 3.**
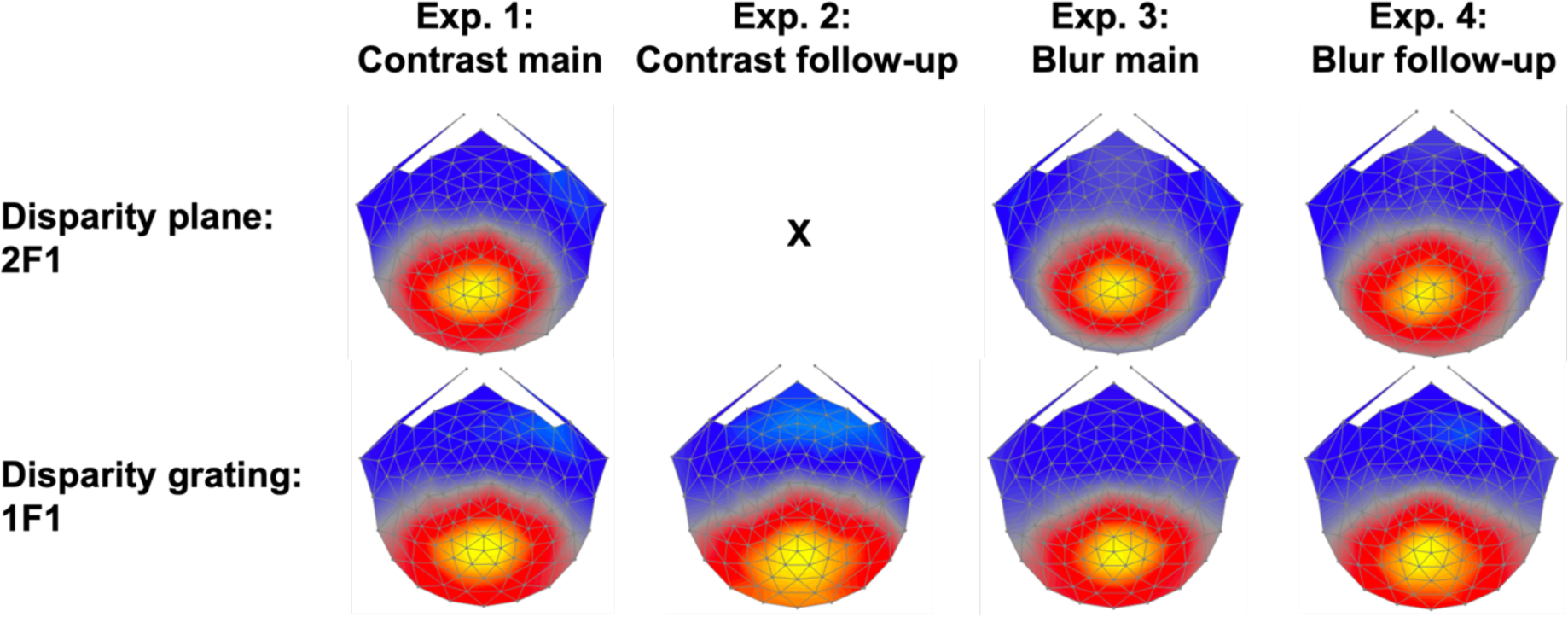
Topographies of RC1 across Experiments and response type. The top row shows RC1 topographies for disparity plane responses learned over the 2F1 response harmonic. The second row shows RC1 topographies for disparity grating responses learned over the 1F1 response harmonic. Response topographies are broadly similar over the four experiments: Experiment 1: Disparity response functions for plane and 0.7 cpd grating conditions, with contrast varying between 1.25 and 40%. Experiment 2: Disparity response functions for gratings of 0.7 and 1.2 cpd gratings with contrast varying between 1.25 and 40%. Experiment 3: Disparity response functions for disparity plane and a 0.5 cpd disparity grating over blur levels between 0 and 4.1 arcmin. Experiment 4: As in Experiment 3, but over an expanded range of blur levels. Spatial filters were learned via Reliable Component Analysis and electrode-level data were projected through each set of weights in further analysis.

### Experiment 1: Contrast response functions for disparity plane modulation

To assess how half-image contrast affects the disparity-specific response to a stimulus that contains primarily absolute disparity, we measured SSVEP amplitude as a function of stimulus disparity for the disparity modulating plane condition. The data are shown in Fig. 4A for the 1F1 component and in Fig. 4B for the 2F1 component. The response to the plane stimulus was larger overall at the second harmonic (2F1; Fig. 4B) than for the first harmonic (1F1; Fig. 4A), as reported previously (Kaestner et al., 2022). Also as has been previously reported (Norcia et al., 1985a; Wesemann et al., 1987; Kaestner et al., 2022), 2F1 amplitude to the modulation of a disparity plane is a linear function of log disparity. As expected, the highest contrast conditions generate the largest amplitude responses, whilst signal in the two lowest contrast conditions did not rise above the experimental noise level (gray band) for either 1F1 2F1 (lightest blue curves in Fig. 4A,B). The high contrast 2F1 response rises out of the noise at around 1.4 arcmin.

**Figure. 4.**
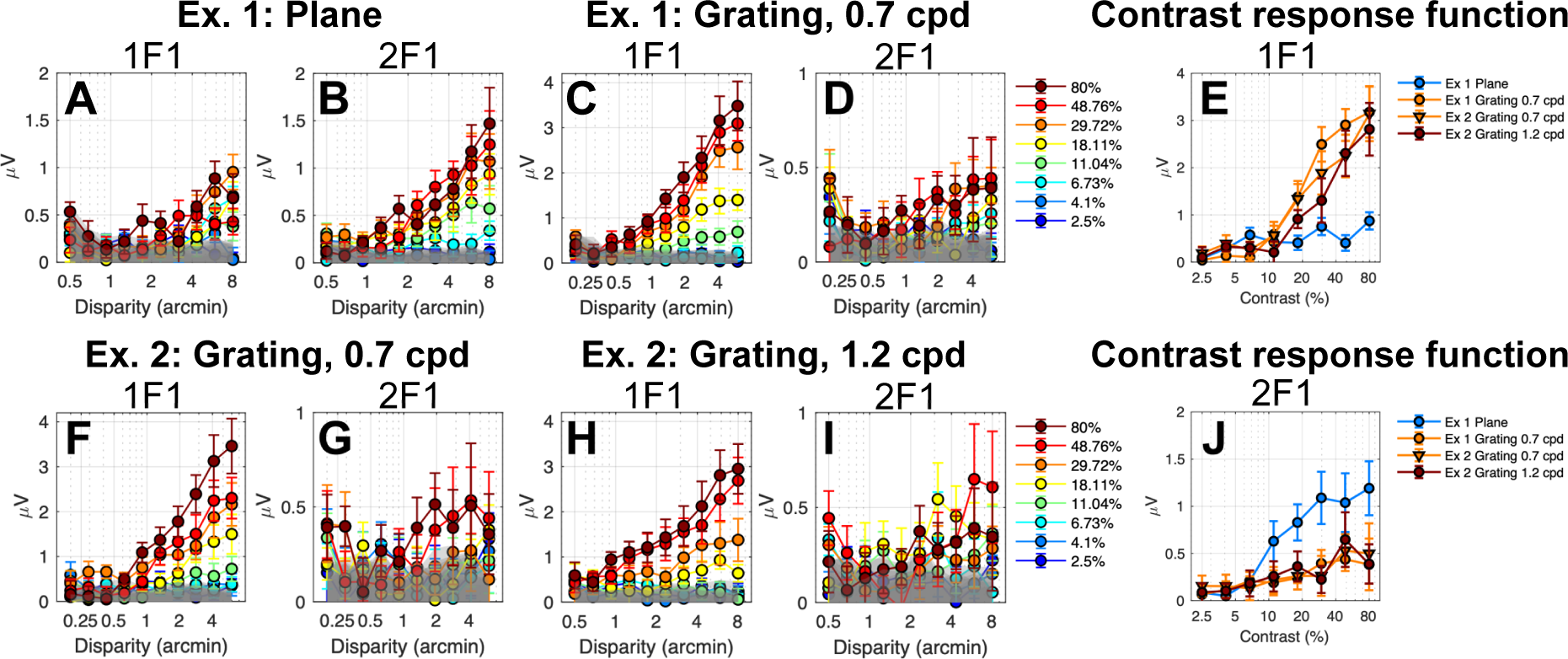
Disparity response functions for plane (A, B) and grating (C,D, F-I) stimuli as a function of contrast (legend) for 1F1 and 2F1. E) Contrast response function derived from responses measured at 6 arcmin of disparity at 1F1. J) As in E), but for the 2F1 response component. Data are show over both Experiments 1 (Ex. 1) and 2 (Ex. 2).

To provide a contrast response function for the disparity plane condition (the nominal absolute disparity condition), we re-plotted response amplitude over contrast levels at a representative supra-threshold disparity (6 arcmin) that was on the linear portion of the response functions for both 1F1 and 2F1. These functions are plotted in Fig. 4E and 4J. The response at 2F1 rises out of the noise level above about 5% contrast and then saturates at 25% contrast (Fig. 4J, blue curve). The response at 1F1 for the disparity plane is close to the noise level and increases only modestly at higher contrasts Fig. 4E, blue curve).

### Experiment 1: Contrast response functions for 0.7 cpd disparity grating modulation

The 1F1 component of the response to a 0.7 cpd disparity grating is ∼3 times larger than the 1F1 response for the disparity plane in both Experiment 1 (Fig. 4C vs Fig. 4A) and Experiment 2 (Fig. 4F), consistent with a contribution from the relative disparity information that is present in the grating, but not the plane stimulus (Kaestner et al., 2022). As was the case for 2F1, 1F1 amplitude for an 0.7 cpd grating is a linear function of log disparity for disparity grating modulation and the highest contrast again produced the largest response. The 1F1 response to the 0.7 cpd grating rises out of the noise at around 0.5 to 1 arcmin, *eg.* at a lower disparity than needed to generate a response to the disparity plane condition (note difference in abscissae of Figs. 4A and 4B vs 4C and 4D).

Replotting the 0.7 cpd grating 1F1 data from Exp. 1 (Fig. 4C) and Exp. 2 (Fig. 4F) as a function of contrast, it can again be seen that the 0.7 cpd grating 1F1 response is much larger than the 1F1 response to the plane (Fig 4E, orange curves vs blue curve). As noted above, the plane 1F1 response is only weakly dependent on contrast above 6%, but the 0.7 cpd grating response increases by a factor of ∼8 in both Experiments 1 and 2. Because of this differential contrast dependence, we suggest that the 1F1 signals generated by the plane and grating arise from different mechanisms, with the grating response reflecting a large contribution from the presence of relative disparity in the stimulus. The contrast dependence seen at 2F1 also differs between the plane and 0.7 cpd grating responses. Here the 2F1 response (Fig. 4J, blue curve) is larger for the plane than the grating (orange curves), rather than being the other way around as in the 1F1 response.

### Experiment 2: Probing the high spatial frequency limb of the Disparity Sensitivity Function

If we consider the plane stimulus to be a very low spatial frequency cyclopean pattern that anchors the low spatial frequency end of the disparity sensitivity function (DSF), the observation that plane and 0.7 cpd grating responses are dominated by different response harmonics suggests that the DSF has at least two underlying spatial channels. The next experiment was designed to test whether there is possibly another cyclopean channel above the mid-range of cyclopean spatial frequencies, as suggested by prior psychophysical work (Peterzell et al., 2017; Reynaud and Hess, 2017). To make this test, we recorded from a 1.2 cpd disparity grating which is on the ascending limb of the DSF (Kaestner et al., 2022), along with a cyclopean grating of 0.7 cpd. Different response slopes would suggest the presence of another disparity channel.

The disparity response functions parametric in contrast for the 1.2 cpd grating are shown in panels Fig. 4H and 4I for 1F1 and 1F2 respectively. The 1F1 responses rise above the noise level by 1 arcmin. The summary contrast response functions for the 1.2 cpd grating are shown in Fig. 4E as the red curve. Responses at 1F1 for the 1.2 cpd grating are rightward shifted on the contrast axis compared to 1F1 responses for 0.7 cpd measured in either Exp.1 (orange circles) or Exp. 2 (orange triangles) as shown in Fig. 4E. The slopes of the disparity response functions for 0.7 and 1.2 cpd gratings are similar. This pattern is consistent with 0.7 and 1.2 cpd gratings activating a common cyclopean channel. It is possible that a different choice of frequencies, *e.g*. lower than 0.7 cpd, higher than 1.2 cpd or both would have shown stronger evidence in the form of different response slopes, for example. The 2F1 responses are the same for 0.7 and 1.2 cpd disparity gratings (Fig. 4J) as reported previously for high contrast DRDS (Kaestner et al., 2022). Here we see that this independence of spatial frequency holds over a wide range of contrast.

### Contrast Response Function summary for plane and grating stimuli

Response amplitudes differ between plane and grating responses at the response harmonics where the signal is largest and most easily measured for the two conditions. To better compare their relative contrast sensitivities, we normalized each function to its peak response in Figure 5, left and middle panels. Each of the functions has a similar slope, but they differ in their position along the contrast axis. The 2F1 response to the plane is located to the left of the 1F1 response function for the 0.7 cpd gratings (orange curve vs purple curves) and the 1.2 cpd grating (green curve). We estimated contrast thresholds for these functions by linear extrapolation to zero amplitude (Campbell and Maffei, 1970; Norcia et al., 1989). The linear fits to the response functions are shown in the middle panel of Fig. 5. The contrast threshold of the dominant response component for the plane was lower (4.4% +/− 0.5%) than for either of the 0.7 cpd 1F1 responses (6.3% +/− 1.1%, Exp. 1; 7.1% +/− 1.7%, Exp. 2) or the 1.2 cpd 1F1 response (10.4% +/− 2.2%) as can be seen in Fig. 5C. Paired-samples *t*-tests run on conditions within each experiment revealed that both pairs of threshold estimates were significantly different after Bonferroni correction for multiple comparisons (Exp 1: 0 cpd plane contrast threshold was significantly lower than the 0.7 cpd grating contrast threshold, T(7) = – 1.27, *p* < .001; Exp 2: 0.7 cpd grating contrast threshold was significantly lower than 1.2 cpd grating contrast threshold, T(7) = –8.41, *p* < .001). The 2F1 component for the plane thus has the lowest contrast threshold (most leftward shifted function) of any of the disparity-specific responses, a novel finding. The 2F1 responses to the grating also appear to have a low contrast threshold, but the shallow slope and low amplitudes of these responses make it difficult to accurately extrapolate a threshold, as shallow functions are biased due to greater inflation of the response by mixing with additive EEG noise than are steeper sloped functions (Norcia et al., 1989).

**Figure 5.**
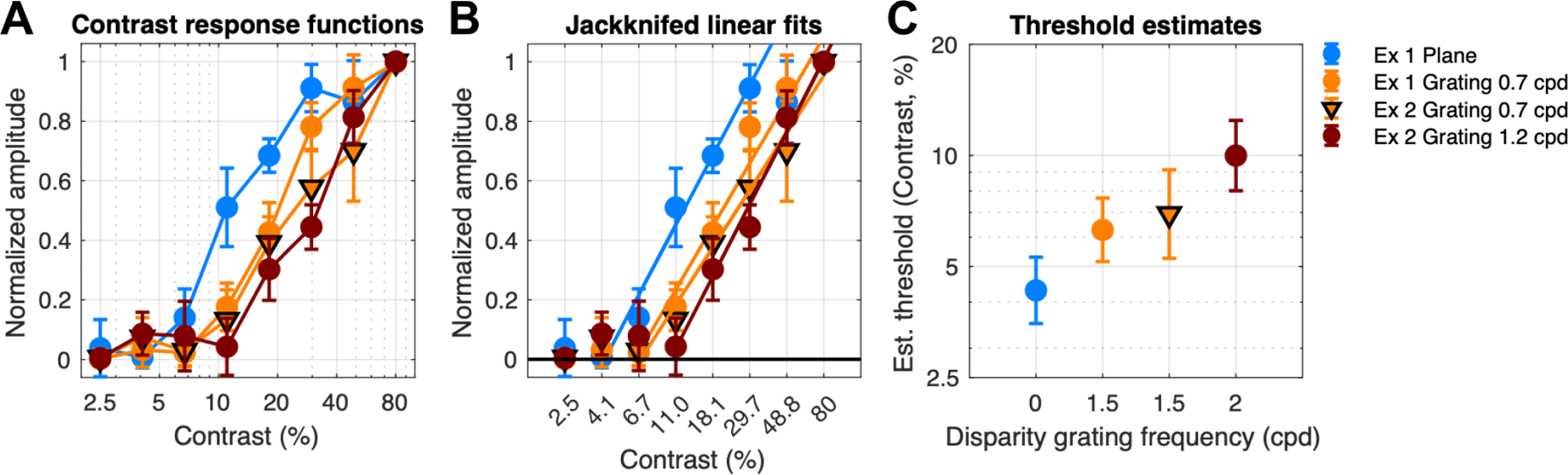
Contrast response functions derived at 6 arcmin from across spatial frequency conditions. Data for the 2F1 response to disparity plane modulation is shown in blue (Exp. 1, Fig. 4B). Data at 1F1 from 0.7 cpd disparity grating modulation are shown in orange (Exp.1, Fig. 4C; Exp. 2; fig 4F). Data at 1F1 from 1.2 cpd grating modulation (Exp. 2, Fig. 4H) is shown in red. Contrast threshold is lowest for the 2F1 response driven by disparity plane modulation, followed by 1F1 thresholds at 0.7 and 1.2 cpd.

### Differential blur sensitivity for plane and grating responses

DRDS patterns are spatially broadband and disparity mechanisms could, in principle, use only a portion of the visible bandwidth available at the input to binocular processing. We thus measured disparity response functions at different levels of blur in two experiments to assess use of high spatial frequency information by the mechanisms generating 1F1 and 2F1 disparity responses.

In experiments 3 and 4, we recorded responses from a disparity plane, a 0.5 cpd disparity grating and a 1.2 cpd grating, using different ranges of blur in the two experiments. Figure 6 provides a composite summary of the disparity response functions for the dominant response harmonics – 2F1 for the disparity plane and 1F1 for the disparity gratings.

A blur of 1.4 arcmin has no measurable effect on either the 2F1 response to the plane (Fig. 6A), the 1F1 response to the 0.5 cpd grating (Fig. 6B) or to the 1.2 cpd grating (Fig. 6C). A blur of 4.1 arcmin shifts the grating response functions to the right for the 0.5 cpd grating (Fig. 6B) and more so for the 1.2 cpd grating (Fig. 6C). This level of blur lowers the slope, but not the apparent threshold for the 2F1 response to the plane (Fig. 6A).

**Figure 6.**
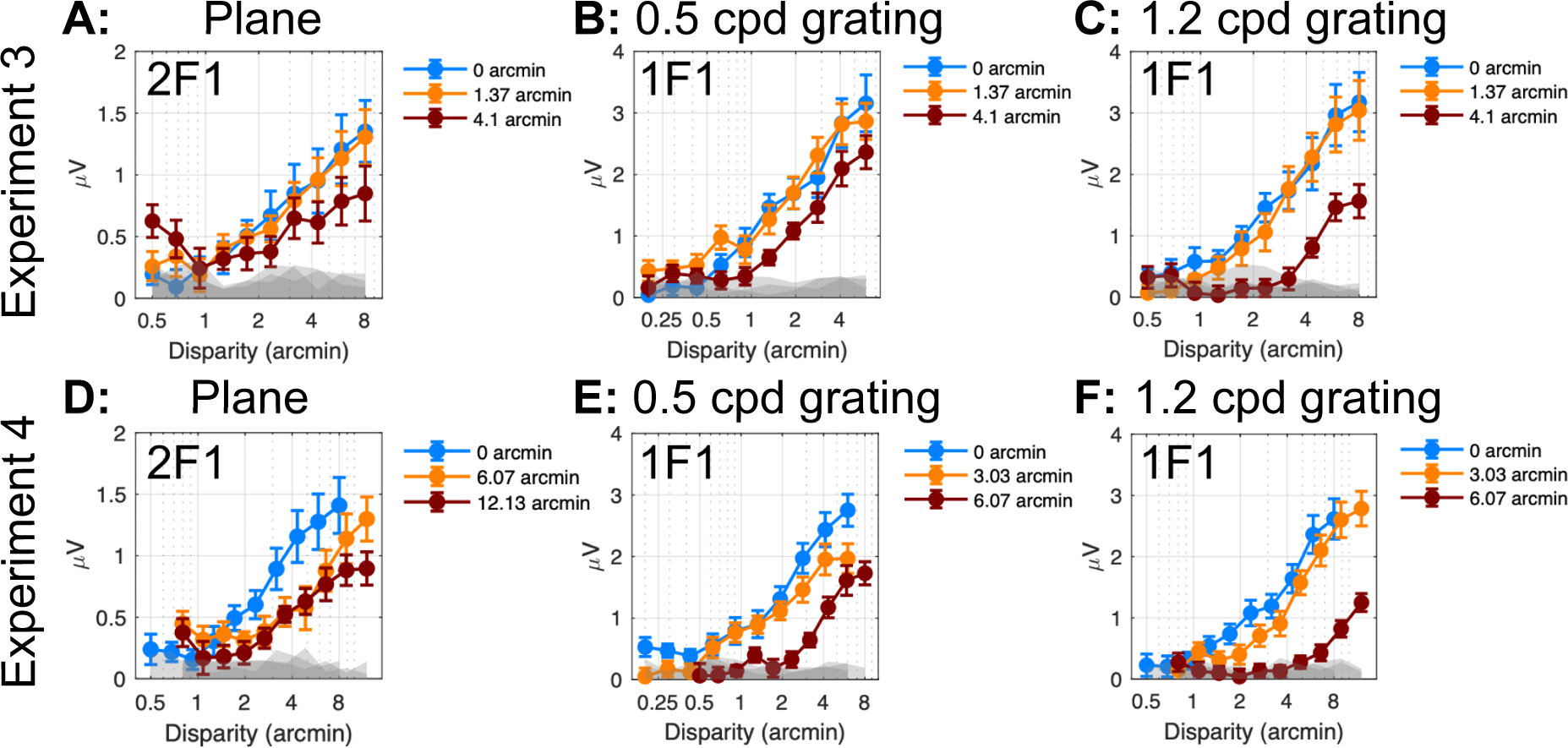
RC1 disparity response functions vs magnitude of imposed stimulus blur. **A:** 2F1 response vs disparity for changing disparity modulation the plane stimulus, Experiment 3. **B:** 1F1 response vs disparity for changing disparity modulation the 0.5 cpd grating stimulus, Experiment 3. **C**: Response vs disparity for changing disparity modulation of a 1.2 cpd grating stimulus (Experiment 3). D) as in A for Experiment 4 E) as in B for Experiment 4. F) as in C for Experiment 4. Gray bands indicate the EEG noise level at adjacent frequencies. Error bars are 1 SEM.

Experiment 2 used larger ranges of blur for both plane and grating stimuli. At 1F1, a blur of 6 arcmin has a greater effect than 4 arcmin of blur on the response to the 0.5 cpd grating (Fig. 6E vs Fig. 6B, note difference in x-axis scale) and similarly an even larger effect on the response to the 1.2 cpd grating (Fig. 6F vs 6C). For the 2F1 response function for the plane condition, a blur of 6 arcmin had a similar effect as a 4.1 arcmin blur in Exp.1. The 12 arcmin blur only affected amplitude at the largest disparities.

To better characterize the effects of blur on disparity processing as manifested by the different response harmonics (2F1 for the plane and 1F1 for the gratings), we estimated disparity thresholds by extrapolating the disparity response functions to zero amplitude. Disparity thresholds were estimated for the three disparity spatial frequency conditions at different blur levels measured over the two experiments and are shown in Figure 7. We then fit polynomial functions to the data for descriptive purposes. Thresholds were constant for the 2F1 plane response up to 12 arcmin of Gaussian blur. By contrast 0.5 cpd grating thresholds were elevated at 4 arcmin of blur and 1.2 cpd grating thresholds were elevated by 2.5 arcmin of blur. The different blur sensitivity between plane and grating response suggest that the former derives from a spatially coarse mechanisms, while the latter derives from finer spatial mechanisms, especially at 1.2 cpd.

**Figure 7.**
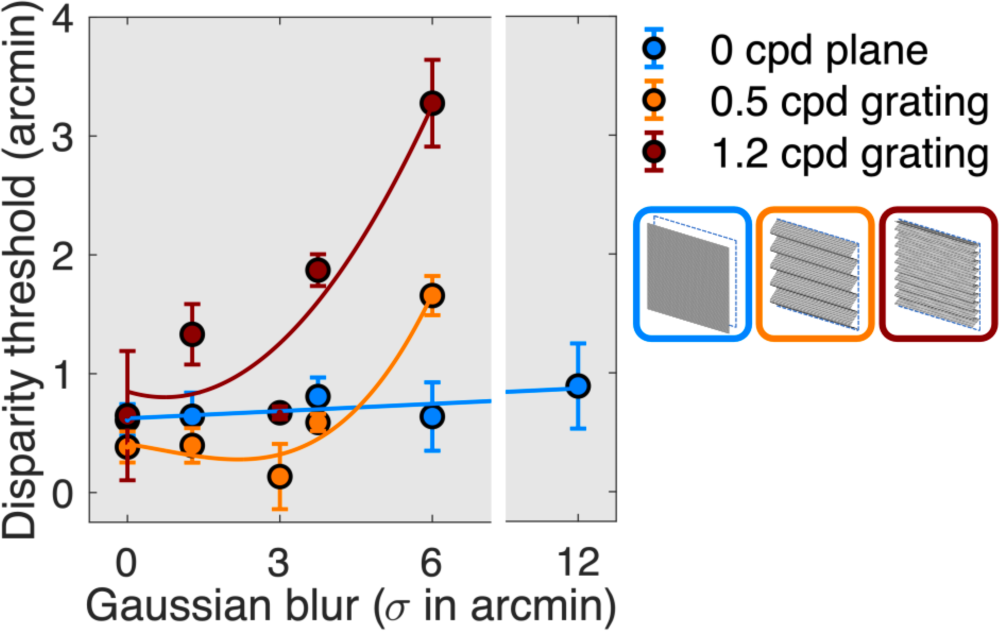
RC1 disparity thresholds as a function of blur for plane (blue), 0.5 cpd grating (orange) and 1.2 cpd grating (red). Data are pooled over the first and second blur experiments.

### Experiment 5: Contrast and blur sensitivity for the half-images

In previous sections we described the contrast sensitivity of different disparity mechanisms. For these disparity-specific mechanism to be active, the monocular half-images need to be visible. Here, we wished to determine the relationship between the contrast thresholds for different disparity mechanisms and the contrast sensitivity of the mechanisms that feed them. To accomplish this, we measured contrast and blur thresholds for the DRDS half-images by modulating the contrast or blurriness of the half-images at 2 Hz (F1). In the former case the images alternated at 2 Hz between 0 contrast and 10 levels of increasing contrast and in the latter the modulation was between unblurred and blurred images with progressively more blur applied in 10 equal log steps. These two measurements allow us to determine if the disparity system is using the full bandwidth available at the input to binocular processing, or only part of it.

Figure 8 shows response topographies and response functions for the 1F1 and 2F1 response components driven by contrast modulation (panel A) and by blur modulation (panel B). Recall that these response components are not disparity-related, as are the responses discussed previously and there was in fact no disparity between the DRDS half-images.

**Figure 8.**
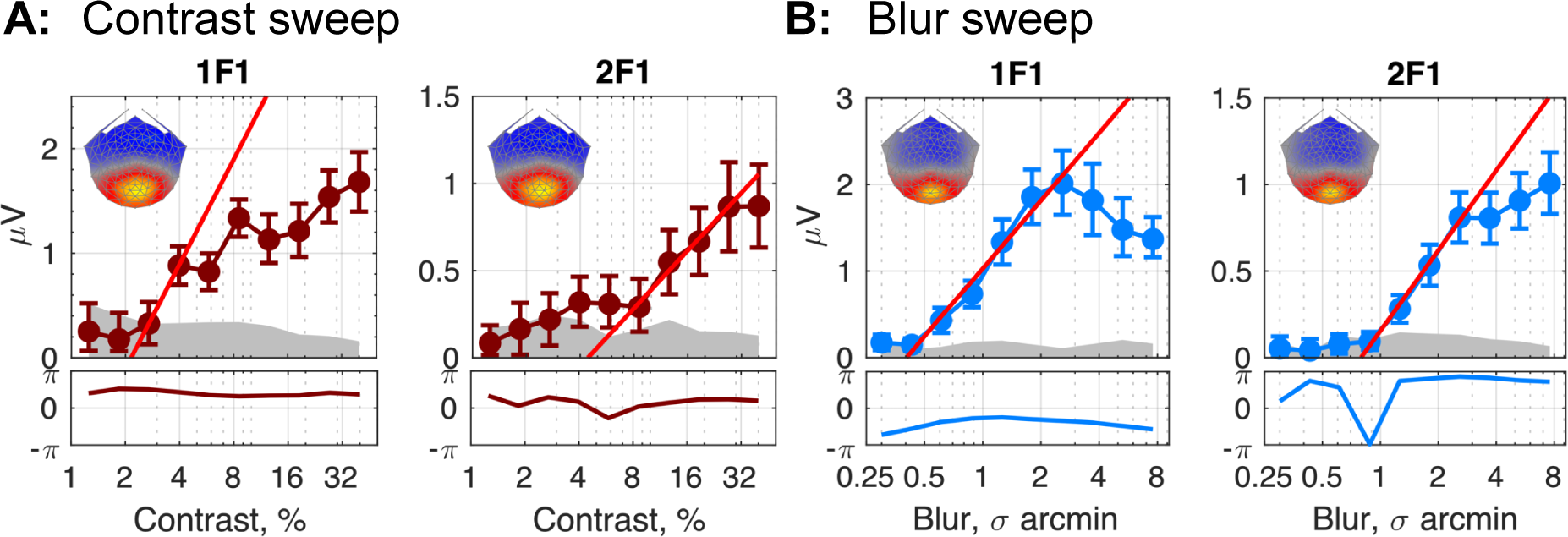
Response functions in RC1 (spatial map insert in each figure) for the first two harmonics of the neural response to a stimulus increasing in contrast (A) or blur (B). Linear fits (bright red line in all plots) reveal the smallest amount of contrast or blur required to elicit a differential response between a given level of contrast or blur, and an unmanipulated image (100% contrast or 0 blur). Generally, this threshold value is smaller for the 1F1 than for the 2F1 response.

The response topographies are each centered over the occipital pole (spatial map inserts in each plot in Figure 8) and resemble those for the disparity-specific responses shown in Figure 1. The contrast response is an approximately linear function of log contrast, as has been previously reported for grating stimuli (Campbell and Maffei, 1970). The 1F1 response has a lower threshold of 2.15 % Michelson contrast than the 2F1 response at 4.42 % Michelson contrast. The blur response function at 1F1 increases monotonically with a neural threshold starting around 0.40 arcmin of blur, over-saturating starting around 3 arcmin. The 2F1 response has a higher initial threshold (0.79 arcmin of blur) and does not saturate. The better of these two thresholds (0.4 arcmin of blur) is in the range of several psychophysical studies of blur detection (Watson and Ahumada, 2011). Together we consider these functions to define the peak contrast sensitivity and high spatial frequency limits of the monocular inputs to disparity processing.

## DISCUSSION

Our goal is to describe the contrast sensitivity of multiple disparity sub-systems and to relate them to the contrast sensitivity of their monocular inputs. That is, we are interested in putting limits on a disparity response component’s “use” of the contrast or spatial frequency content of DRDS half images. The present work and previous work (Chen et al., 2022; Kaestner et al., 2022) suggests that stimuli containing absolute and relative disparity generate responses in at least two disparity sub-systems whose activity is largely reflected by different response harmonics of the changing disparity SSVEP, with 1F1 responses being prominent for stimuli containing relative disparity and 2F1 responses predominating for stimuli that contain only absolute disparity. By manipulating stimulus contrast and blur, we estimated the underlying contrast sensitivity functions for these different response components and by inference, the sensitivity of absolute and relative disparity mechanisms. If a component shows no effect of contrast or blur above or below a certain level, then those contrasts or spatial frequencies are considered to not be used by the mechanisms that drive the particular response component.

### Contrast sensitivity for half-images

We estimated the contrast sensitivity function *(e.g.* contrast threshold as a function of spatial frequency) underlying the processing of the half-images themselves – independent of disparity – from our contrast and blur modulation measurements. The contrast modulation experiment determines the minimal amount of half-image contrast that evokes a response and is a detection threshold. The blur modulation experiment determines the minimal amount of blur that produces a response that differs from that generated by unblurred half-images. It is thus a spatial frequency discrimination threshold in the broad sense. The derived contrast sensitivity function reflects contributions from monocular inputs to cortex as well as binocular cortical responses. The latter could be either tuned for disparity or not. This “front-end” function is expected to limit downstream disparity sensitivity. In Figure 9A, we construct limits for this function by placing a horizontal line at the contrast sensitivity estimated from the contrast sweep data in Fig. 8C and a vertical line at the spatial frequency limit estimated from the blur sweep of Fig. 8A. We then assume a low-pass spatial filter function based on the fact the DRDS image content itself is updating at a rapid rate (Robson, 1966). All other sensitivity functions for disparity-specific mechanisms should lie within these bounds and this is the case empirically. Peak half-image contrast sensitivity is ∼50 and we set the spatial frequency cut-off (somewhat arbitrarily) at 30 cpd, referencing the other functions to this value by their multiples of the blur threshold for the half-images (0.4 arcmin, Figure 8).

**Figure 9.**
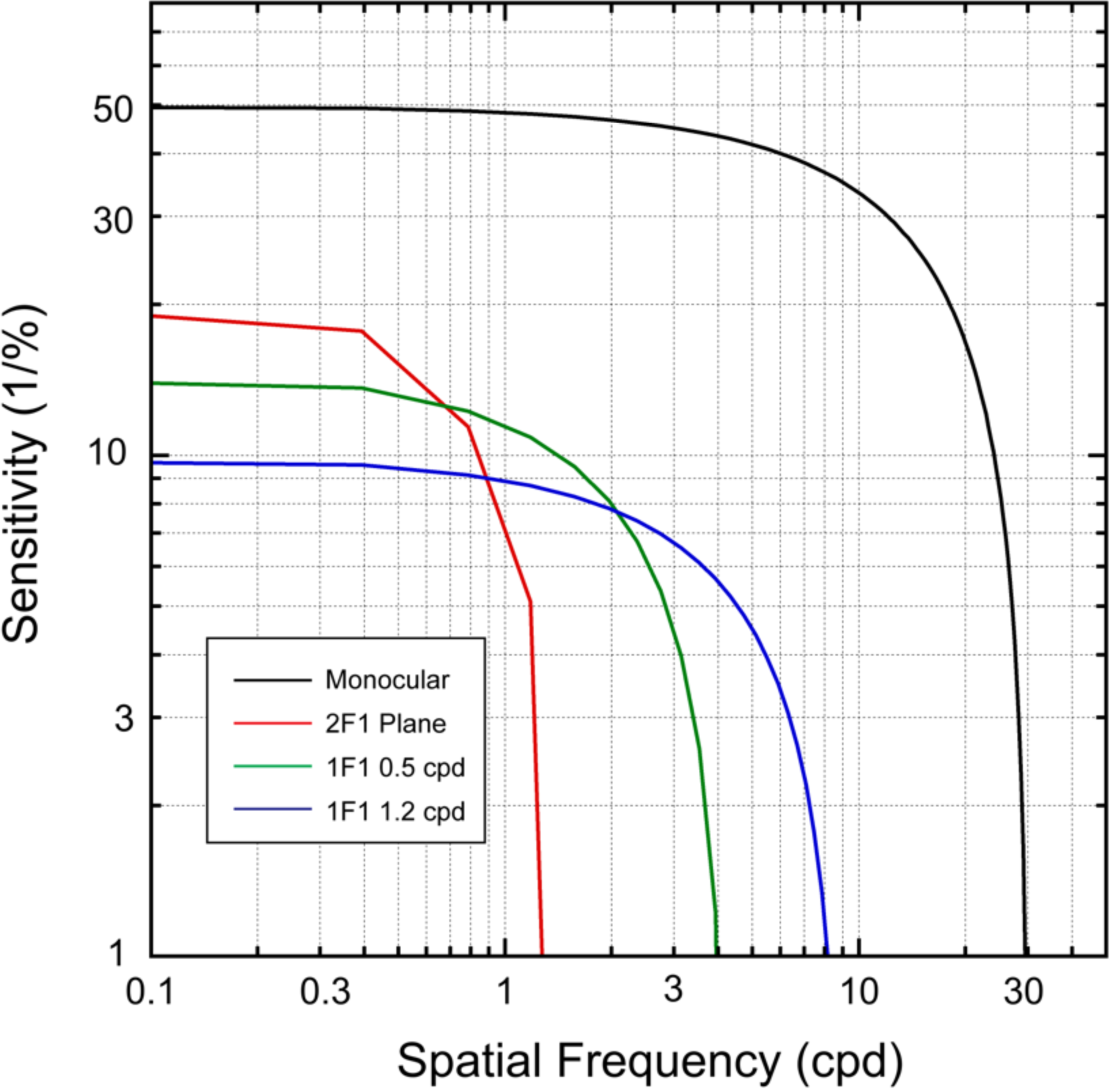
Summary of inferred contrast sensitivity functions (CSF) for different response components. Black curve: CSF for the monocular half-images derived from Experiment 5, 1F1. Red curve: CSF for the 2F1 responses to disparity modulation of a plane. Contrast sensitivity is lower than for the half-images, but higher than those for the grating responses. The high spatial frequency limit is the lowest measured. Green: CSF for 0.5-/0.7 cpd disparity gratings. Peak contrast sensitivity is intermediate between the plane and 1.c cpd gratings, as is its spatial resolution. Blue: The CSF for the 1.2 cpd disparity grating has the lowest peak sensitivity, but the highest resolution of the three disparity-dependent CSFs.

### Contrast sensitivity function for the disparity plane (2F1)

We estimate the contrast sensitivity function underlying the dominant response to the disparity plane (2F1) from the measurements of amplitude *vs* contrast shown in Fig. 5 and from the blur sweep functions shown in Fig. 7. Here the estimated contrast sensitivity for the 2F1 component of the disparity plane response to be ∼20 or about a factor of ∼2.5 lower than the sensitivity measured for the half-images. The threshold of the disparity 2F1 component is unaffected by the largest blur we used, 12 arcmin of blur (see Fig. 7). This response is highly resistant to blur, suggesting that it is generated by coarse disparity mechanisms, but we can only place an upper limit, here of ∼ 1cpd.

The 2F1 response has previously been associated with transient disparity processing (Chen et al., 2022), based on its temporal waveform and similarity to previous results for contrast evoked VEPs (McKeefry et al., 1996). Consistent with this, the temporal frequency tuning of the SSVEP second harmonic to a changing disparity plane is bandpass, peaking at around 3 Hz (Norcia and Tyler, 1984). Our previous work has shown that the response at 2F1 is not tuned for cyclopean spatial frequency (Kaestner et al., 2022) and is larger for disparity planes than gratings as can also be seen in the present results. Together, these results suggest that the 2F1 recorded in response to changing disparity plane reflects – to a substantial degree – the transient processing of absolute disparity by spatially coarse mechanisms.

### Contrast sensitivity functions for disparity gratings (1F1)

Peak contrast sensitivity for the 0.7 cpd (∼15) and especially the 1.2 cpd gratings (∼10) are lower than that for the disparity plane (∼50). This suggests that contrast sensitivity is higher for absolute than for relative disparities – a novel finding. The derived high spatial frequency limits from the blur manipulation for 0.5 (∼4 cpd) and 1.2 cpd (∼8 cpd), are higher than measured for the disparity plane at 2F1 (∼1 cpd). This indicates that the mechanisms underlying 1F1 for disparity gratings are finer spatially than the mechanism underlying the dominant 2F1 response to the plane/absolute disparity condition. Note that the bounding line approach we use to infer the underlying contrast sensitivity functions assumes a low-pass characteristic. It is possible that the tuning for the grating channels could be bandpass instead of low-pass.

We have previously associated the 1F1 response to changing disparity with temporally sustained disparity processing (Chen et al., 2022; Kaestner et al., 2022). Moreover, the 1F1 response here and in our previous work is much larger for disparity gratings that contain relative disparities than it is for a modulating disparity plane which does not (Chen et al., 2022; Kaestner et al., 2022). Explicit manipulation of the availability of references also strongly modulates the 1F1 response (Cottereau et al., 2011, 2012b; Cottereau et al., 2012a; Kaestner et al., 2022) with 1F1 being larger in the presence of a disparity reference, also indicating that it reflects the processing of relative disparity. Taken together, these results suggest that 1F1 generated from the onset/offset of spatial patterns of disparity reflects, in large part, the sustained processing of relative disparities by spatially fine mechanisms.

### Specificity of 1F1 and 2F1 responses

To what extent does the existing data support a strict association of 1F1 responses with the sustained coding of relative disparity and 2F1 response with the transient coding of absolute disparity? Because the SSVEP is a mass response arising from large populations of neurons in different cortical areas, such a strict dissociation without considering the stimulus conditions used seems unlikely even though the use of DRDS stimulation restricts the relevant population to one that is sensitive to disparity. Within this broader population of disparity sensitive cells, different sub-populations could contribute to the different response components and their contributions could depend on the stimulus used. Populations contributing to the 1F1 response could indeed include cells that are tuned for relative disparity if that is present in the stimulus. However other activity at the population level not specific to relative disparity could also contribute and be apparent, say, in the 1F1 response for the disparity plane, a nominal absolute disparity stimulus. Prior modelling and experimental work has suggested that both first and second harmonic responses can arise from a population of cells tuned only for absolute disparity if the population is comprised of a mixture of cells narrowly tuned to zero disparity and other cells tuned for non-zero absolute disparity (Cottereau et al., 2011). Alternatively, population-levels responses could reflect asymmetries between motion towards and motion away from the observer, given that we present changing disparities that could be coded by cells tuned for direction of motion in depth. A population-level asymmetry in 3D motion could thus manifest at 1F1. Responses at 1F1 could reflect increasing amounts of decorrelation as disparity increases in a population tuned to zero disparity. Finally, residual relative disparity responses may be present due to incomplete elimination of references in the disparity plane condition. This is unlikely because the 1F1 response to a disparity plane is insensitive to blur (data not shown) rather than being sensitive to blur as is the 1F1 to the disparity grating. Importantly, explicit manipulations of references that do not modify motion in depth or the levels of correlation/decorrelation have large effects on 1F1 (Cottereau et al., 2011, 2012b; Cottereau et al., 2012a; Kaestner et al., 2022), indicating that the 1F1 response is strongly influenced by relative disparities, when they are present.

Similar considerations apply to the 2F1 response. Transient response mechanisms will generate second harmonic responses (McKeefry et al., 1996). These responses are largest for the disparity plane but are also present for the disparity gratings. Disparity gratings have both absolute and relative disparities in them and the 2F1 response to gratings may receive a contribution from transient activity of a population of absolute disparity cells. It is also likely that cells tuned to relative disparity are not purely sustained, but rather display a (small in adults) degree of transiency in their response. Finally, 2F1 response could reflect some unspecified form of non-linear activity in a population of cells tuned for relative disparity. These responses are small and more difficult to measure, and further work is needed to clarify the mechanisms that underlie their generation.

Rather than considering the 1F1 and 2F1 responses in isolation, a stronger case for their linkage to separate disparity-selective processes comes from associating them with stimuli that optimize them, e.g. 1F1 from a disparity grating or other stimulus that contains relative disparity and 2F1 from a stimulus that does not contain relative disparity.

### Relationship to previous studies

To our knowledge there are no single-unit or functional MRI measurements of the contrast required to generate a disparity-specific response or how the contrast sensitivity of disparity tuned cells relates to that of other cells that are not. We find that disparity sensitive cells are less sensitive to contrast than other cells that are not. The disparity system thus only appears to use a portion of the monocularly visible image bandwidth.

One study using the VEP has measured a contrast response function for alternations between correlated and anti-correlated DRDS patterns with coarse pattern elements (Marko et al., 2009). A VEP response to correlated/anticorrelated transitions is clearly a binocular response and may also arise from cells that have some degree of disparity tuning (Poggio et al., 1988). Marko and co-workers found that response amplitude was nearly constant between 5.5 and 80% contrast, suggesting a mechanism with high contrast sensitivity like the one we observe for the disparity plane responses at 2F1.

Measurements of vergence eye movements have suggested that the relevant disparity inputs are spatially coarse (Westheimer and Mitchell, 1969; Jones and Kerr, 1972; Edwards et al., 1998; Pope et al., 1999; Sheliga et al., 2006), consistent with our finding of high resistance to blur of the disparity response to changes in absolute disparity – the presumptive disparity input to vergence. Disparity vergence responses to DRDS have high contrast sensitivity and saturate at moderate contrast levels (Stevenson et al., 1994), consistent with our results for 2F1 of the disparity plane response. Finally, disparity vergence can be driven by coarse stimuli, even ones that cannot be binocularly fused (Westheimer and Mitchell, 1969). The tuning of the inputs to our 2F1 response to changing absolute disparity thus shares properties of spatial coarseness, high contrast sensitivity with the inputs to disparity vergence.

It is well-known that stereo acuity, a relative disparity judgement, is strongly dependent on image contrast (Halpern and Blake, 1988; Legge and Gu, 1989; Cormack et al., 1991) and blur (Brooks et al., 1996; Banks et al., 2004; Atchison et al., 2020a; Atchison et al., 2020b). These findings are corroborated by our measurement of the 1F1 grating response, which was sensitive to both contrast and blur.

Prior psychophysical work that has examined the relationship between the luminance spatial frequency of the stereo half-images and sensitivity to different spatial frequencies of disparity modulation (corrugation frequency) is relevant to our blur experiments. An early study (Pulliam, 1981) imposed sinusoidal disparity modulations of differing spatial frequencies on vertical luminance gratings of different spatial frequencies and concluded that channels tuned for fine disparity modulation frequencies were tuned for high luminance spatial frequencies and vice versa. Similar conclusions were reached by (Hess et al., 1999). Other work (Lee and Rogers, 1997), by contrast, suggested that a single luminance channel tuned to around 4 cpd supported optimal disparity corrugation detection. (Witz and Hess, 2013) reconciled these results by suggesting that the optimal luminance spatial frequency increases with increasing cyclopean spatial frequency above 1 cpd, but is constant at ∼3 cpd for cyclopean spatial frequencies lower than 1 cpd. Our blur results are qualitatively consistent with an association of high luminance spatial frequency with high disparity spatial frequency and vice versa. But Witz and Hess’s cutoff at 1 cpd between regimes suggests that there would not be the large difference we see in blur sensitivity between 0.7 cpd and the disparity plane condition. There are multiple differences between our paradigm and that of Witz and Hess (2013) that may alter this relationship, e.g. we used changing disparities instead of static disparities and our outcome measure was physiological rather than perceptual.

Perceptual sensitivity to disparity corrugations – the DSF – peaks at around 0.5 cpd, as does the sensitivity derived from the 1F1 response (Kaestner et al., 2022). Other psychophysical work has suggested that more than one cyclopean spatial frequency channel underlies the DSF (Peterzell et al., 2017; Reynaud and Hess, 2017). Response differences between the disparity plane and disparity grating conditions suggest the presence of at least two channels underlying the DSF. Disparity plane responses are distinctive in terms of their temporal dynamics, being dominated by 2^nd^ rather than 1^st^ harmonics, as are disparity grating responses. These 2^nd^ harmonic plane responses have lower sensitivity to modification by blur than the 1F1 responses to gratings, but higher contrast sensitivity. Moreover, the 2F1 response to disparity gratings are untuned for cyclopean spatial frequency (Kaestner et al., 2022) and Fig. 4J. Thus, because of these differences in the tunings of 1F1 and 2F1, we have evidence for more than one “channel” between the peak of the DSF and low cyclopean spatial frequencies.

Our results are equivocal on whether there is third channel underlying the high-spatial frequency limb of the DSF. At the high spatial frequency end of the DSF, response functions for 1.2 cpd gratings appear to be laterally shifted on either contrast or blur axes, with no qualitative differences such as a change of slope relative to our lower spatial frequency cyclopean grating responses. Lateral shifts could arise from a reduction of sensitivity of a single channel to higher cyclopean spatial frequencies, or they could come about from the intrinsic sensitivity of a separate channel. The psychophysical channels were derived from a co-variance analysis of psychophysical thresholds. A similar co-variance analysis could be performed on the 1F1 and 2F1 SSVEP data over the entire DSF to determine whether the 1F1 response to gratings shows evidence of multiple channels sensitive to relative disparity.

## Conclusion

Returning to our initial translational goal of validating the SSVEP as a useful index of stereoscopic sensitivity, we can suggest that in normal vision adults, the 1F1 response to a modulating disparity grating provides a neural correlate of spatially fine and temporally sustained relative disparity processing and the 2F1 response to a disparity plane indexes spatially coarse and temporally transient absolute disparity processing. It will be of interest in the future to determine whether these processes are differentially affected by disorders of binocular vision such as strabismus and/or amblyopia and whether they are differentially responsive to treatment.

## Acknowledgements

This research was supported by grant no. EY018875 from the National Eye Institute, National Institutes of Health. The authors would like to thank Vladimir Vildavski and Alexandra Yakovleva for the development of instrumentation used in the experiments.

## Notes

### Competing Interest Statement

The authors have declared no competing interest.

